# Species-specific responses of helminths to temperature and moisture: long-term and multi-scale analyses in a free-living rodent

**DOI:** 10.64898/2026.04.21.719831

**Authors:** Abosede E. Olarewaju, Urszula Zawadzka-Pawlewska, Victoria I. Ayansola, Alyssa Dunn, Anna Rybińska, Anna Bajer, Jerzy Behnke, Mohammed Alsarraf, Dorota Dwużnik-Szarek, Katarzyna Tołkacz, Maciej Grzybek, Jolanta Behnke-Borowczyk, Agnieszka Kloch

**Affiliations:** University of Warsaw, Institute of Ecology, Faculty of Biology, Warsaw, Poland; University of Warsaw, Faculty of Geography and Regional Studies, Warsaw, Poland; Pitzer College, 1050 N. Mills Avenue, Claremont, CA 91711, USA; University of Warsaw, Department of Eco-Epidemiology of Parasitic Diseases, Faculty of Biology, Warsaw, Poland; School of Life Sciences, University of Nottingham, University Park, Nottingham, UK; Institute of Biochemistry and Biophysics, Polish Academy of Sciences, Warsaw, Poland; Department of Tropical Parasitology, Institute of Maritime and Tropical Medicine, Medical University of Gdańsk, Gdynia, Poland; Department of Forest Entomology and Pathology, Faculty of Forestry and Wood Technology, Poznań University of Life Sciences, Poznań, Poland

**Keywords:** Climatic variation, Helminth ecology, Host-parasite interactions, Parasite burden, Wild rodent

## Abstract

Parasite infections in wildlife vary across time and space due to interactions among host biology, ecological processes, and climatic variability. Under ongoing climate change, understanding how temperature, precipitation, or humidity influences parasite dynamics is important for predicting shifts in infection patterns and host-parasite interactions. Here, we examine how variation in climatic conditions is associated with helminth infections in a free-living rodent, the bank vole (*Myodes glareolus*), across 17 years and multiple spatial scales. Using zero-inflated generalised linear models, we quantified the effects of climatic variables on individual parasite burden. Climatic conditions (temperature and humidity or precipitation) affected helminth infections across all analysed scales, though the strength and direction of these effects differed among parasite species and between temporal and spatial scales. In the temporal dataset, parasite load was associated with seasonal variation in weather conditions, whereas in the spatial datasets, infection levels were linked to yearly average climatic variables. The differences reflect species-specific parasites’ life histories and transmission strategies. Our findings highlight the importance of analysing individual parasite species rather than overall parasite load or aggregated infection indices when assessing the impacts of climatic variation on host-parasite dynamics.

## 1. Introduction

Parasite communities exhibit significant variation in their distribution and abundance across temporal and spatial scales, shaped by complex interactions among host-related, ecological, and environmental factors. Among these, climatic variables such as temperature, precipitation, and seasonality play an important role in influencing parasite dynamics, particularly at small geographic scales, where microclimatic conditions can create pronounced heterogeneity in parasite communities [1, 2]. Understanding how these climatic factors drive changes in parasite communities over time and space is critical for predicting the impacts of climate change on host-parasite interactions and disease dynamics. Due to the crucial role of parasites in shaping dynamics of their host populations, such predictions seem essential not only for understanding parasites alone but also for the species they infect. Yet, except for the parasites of human health importance, there is a scarcity of the studies examining the effect of climate change on pathogen communities.

Macroecological patterns in parasite distribution and abundance are categorised as either universal (predictable and testable) or contingent (context-dependent) [1]. Universal patterns, such as the relationships among parasite abundance, prevalence, and host population dynamics, often apply across broad geographic areas and host species. In contrast, contingent patterns, often highly context dependent, emerge at the community level, where interactions among parasite species and their hosts are complex and often occur under non-equilibrium states [3, 4]. This complexity makes it challenging to formulate general laws governing dynamics in parasite communities. While studies have explored repeatability of parasite community composition across time or space individually, few have integrated both dimensions, especially in terrestrial systems. Consequently, most research has focused on aquatic environments [5, 6], leaving a gap in our understanding of how climatic factors shape parasite communities across geographic scales over time and space.

Climatic factors such as temperature and precipitation directly influence parasite life-history traits, transmission dynamics, and host susceptibility [7, 8, 9]. For example, seasonal fluctuations in temperature and humidity can alter the availability of intermediate hosts or vectors, while long-term climatic changes can shift the geographic ranges of both hosts and parasites [10, 11, 12, 13]. [14] showed that a remotely sensed primary production index predicts variation in parasite burden. In a recent review of the influence of environmental factors associated with climate change (temperature and/or moisture) on terrestrial parasitic nematodes’ life histories and behaviour, [15] identified a large knowledge gap where 66% of hosts examined were cattle or sheep, and the majority of parasites studied were just three species, all infecting livestock. Similarly, [16] in their meta-analysis identified a knowledge gap around the relationship between humidity and parasitism in terrestrial animals, and they reported a scarcity of studies where the infection intensity was studied instead of prevalence.

Majority of studies aimed at characterizing the links between climatic variables (temperature, humidity, precipitation) and parasite load adopt a cross-taxonomic approach, seeking a broader pattern that would be uniform across species. For instance, the latitudinal diversity gradient (LDG) based on the classic theory of higher species diversity at lower latitudes towards the equator, proposes increasing parasite species diversity with decreasing latitude. While this trend is often observed in free-living organisms, parasites are inconsistent in their relationship with latitude. A more general notion is that parasites will exhibit no latitudinal pattern or reverse gradient [reviewed by 17]. [17] found that nematodes followed the traditional LDG, while cestodes and trematodes exhibited reverse or no latitudinal patterns, suggesting that differences in helminths’ adaptability and responses to environmental conditions could explain the different latitudinal patterns observed in the taxa. A related hypothesis is “warmer sicker world” that predicts an increase in parasite prevalence with the rise of global mean temperatures [18, 19]. However, recent meta-analysis of terrestrial animals did not support this hypothesis [16], and the authors reported large variation in the relationship between temperature, precipitation, and parasite prevalence with no overall visible trends.

These inconsistencies emphasise the importance of considering various scales when studying environmental impacts affecting parasite diversity. On a smaller geographic scale, microclimatic variations can create distinct environmental niches, leading to localised differences in parasite community composition and abundance. These small-scale patterns are often overlooked in broader studies but are also crucial for understanding the mechanisms driving parasite dynamics in response to environmental change, and as a consequence, the effect of parasites on the populations of their hosts. In the present study, we aim to fill the knowledge gap indicated by [16] by investigating how spatially and temporally variable climatic conditions affect parasite communities in a single host species using the widespread, pan-European rodent bank vole (*Myodes glareolus*) as an example.

The bank vole (*Myodes* (*=Clethrionoyms*) *glareolus* (Schreber, 1780)) provides an excellent model for studying the impact of climatic factors on parasite communities across time and space. Bank voles have a well-characterised life history and a broad ecological distribution across diverse environmental gradients in Europe and part of Asia [reviewed by 20]. Long-term monitoring of bank vole populations has revealed a well-characterised community of helminths and other parasites, providing valuable longitudinal data for ecological and parasitological analyses [21, 22, 23]. Previous research has demonstrated that parasite communities in bank voles are relatively stable over time and consistent across several geographic regions [22, 23, 24]. For instance, in a long-term rodent survey, [23] observed consistent differences in the prevalence of several nematodes between sites separated by just 10-25 km, suggesting that small-scale environmental gradients and climatic factors play an important role in shaping parasite dynamics. In other rodent and wildlife systems, year-to-year studies have also highlighted the role of host and ecological factors in shaping parasite dynamics.

In the present work, we analysed the relationship between precipitation and temperature and parasite load at three levels: 1) temporal scale, including several capture sessions of 17 years from 1999 to 2016; 2) medium spatial scale, spanning around 800 km (nearly 5° of latitude) across a range of landscapes; and 3) large spatial scale, including data derived from published papers. By comparing these three scales, we aimed to differentiate the effect of climatic factors that are characteristic of a given locality from factors that vary across seasons. We expected that weather conditions would have a major effect on parasite species that stay outside the host for a long time (e.g., eggs deposited in and larvae developing in the soil). We determined if this pattern is consistent over time and space. Finally, we tested whether latitude, a variable linked to several environmental conditions, can explain regional differences in the parasite load.

## 2. Methods

### 2.1. Temporal dataset

The first dataset (henceforth called “temporal”) consisted of data collected across a 17-year time span in a single location in North-Eastern Poland (Urwitałt, 53.7986N, 21.6547E).

This dataset provided a fine scale where we were able to contrast parasite load with local monthly weather conditions to account for temporal variation.

The temporal dataset contained data on 757 bank voles collected from 1999 to 2016 at the Urwitałt site (**Table S1**). Details of the trapping protocol have been fully described before in earlier studies [22, 25, 26, 27, 28]. Briefly, voles were caught live in locally constructed wooden traps arranged in transects. Vole density was estimated as the number of voles captured per 10^4^ trap-hours. The population density varied considerably, from over 220 voles per 10^4^ trap-hours in late summer of the peak year to as low as five voles in May, but the number of sampled voles did not correlate with the population density (r = 0.304, t = 1.278, p-value = 0.2195).

Voles were weighed (±1g), sexed, and euthanised following the European ethical standards (Directive 2010/63/EU) through anaesthesia with isoflurane followed by bleeding from the heart, except for samples from 2022 and 2023, which were followed by cervical dislocation. The voles were dissected; internal organs were extracted, and the intestinal tract examined for intestinal parasites. All helminths were removed, counted, and preserved in 70% ethanol. Species of helminths were identified using standard parasite morphological descriptions provided in [29,30].

Modelling associations between parasite burdens and weather conditions over time is confounded by two factors: seasonal host dynamics [e.g., 31] and seasonal variability in weather conditions typical of the temperate climatic zone. Here, we addressed this issue by comparing infection intensity with monthly weather conditions at the time the hosts were captured. Moreover, to account for the time needed to develop an infection after the parasite’s infective stages had been encountered, we fitted separate models with weather conditions two and four months prior to capturing. The time window was selected so that it corresponded to the average cohort age estimated using lens weight recorded for a subset of animals [32, 33]. In spring, all analysed voles were adults older than 2.5 months old. Adults (>2.5 months old) and young adults (>1.5 months old) constituted 74.5% of individuals captured in late summer and 78.5% of animals captured in autumn.

Monthly climatic variables for the meteorological station Mikołajki (5 km from Urwitałt) were obtained from the publicly available database of the Institute of Meteorology and Water Management - National Research Institute (IMGW-PIB). The data include mean average, mean minimal, and mean maximal monthly temperatures [°C], the minimal monthly temperature at the ground level [°C], and precipitation (as a number of days with rainfall or snowfall per month).

### 2.2. Medium-scale spatial dataset

The medium-scale spatial dataset comprised 287 voles sampled in 16 locations from six geographic regions across an 800 km span (**Figure S1**, **Table S2**) in 2016, 2022, and 2023. All sites were sampled in August/September 2022 or 2023, except for Mazury, which was studied in September 2016. We found striking differences in vole densities between sites. For instance, in 2023, we captured only 9 voles in Góry Świętokrzyskie (Central Poland) and 13 in Beskid Żywiecki (Southern Poland), but in the same field season with the same trapping effort, we collected over 60 animals in Křižanovská vrchovina in the Czech Republic. There is a vast amount of ecological literature examining the reasons and factors influencing population densities, and searching for suitable explanations is beyond the scope of this study. However, to control for the demographic factors, we fitted population density in the models.

Data collection (trapping, animal handling, etc.) followed the same protocol as the temporal dataset.

Climatic variables were downloaded from several databases. Bioclimatic indicators from CMIP5 climate projections were downloaded from the European Centre for Medium-Range Weather Forecasts and downscaled to 1 km^2^ resolution [34]. The dataset consisted of annual records averaged for the 2001-2020 period: mean temperature (BIO 01, obtained in °K transformed to °C) and annual temperature amplitude (BIO 07, obtained in °K transformed to °C). From the ERA5 database [35], we downloaded the monthly averaged soil temperature at a depth of 3.5 cm (in °K transformed to °C), the volumetric soil water layer, and the depth of the snow cover. The data were downloaded for seasons and averaged to obtain a mean value for the period 2001-2020. Precipitation (mm) was obtained from E-OBS daily gridded meteorological data [36], and the average sum of annual precipitation was calculated for the period 2011-2019.

### 2.3. Large-scale spatial dataset

For this part of the analysis, we searched PubMed for *parasite and *glareolus, or *helminth and *glareolus. We included papers published after 1980, since 1980 marks a threshold after which the temperature anomalies have consistently exceeded the 1950-1980 averages [37]. We excluded data from urban habitats, since host-parasite relationships in such habitats may be influenced by different factors than in natural or semi-natural habitats. Furthermore, we excluded data from the British Isles to avoid the confounding effect of the insular climate, and because voles colonised this area much later than continental Europe. We extracted the location, year of sampling, sample size, and the prevalence of each reported nematode from the papers. If several papers described the same location, we used the most recent data. A summary of the reviewed studies is given in **Table S3**, and the references are listed in the Supplementary Materials.

We collected climatic data for each location similarly to the medium-scale spatial dataset. We also included data analysed in the medium-scale spatial dataset with prevalence instead of infection intensity, discarding locations where less than 10 voles were captured.

The data spanned from 39°N in Italy to 68°N in Finland and from 2°E in the Pyrenees to 45°E in Russia, adequately representing the whole range of bank voles (**Figure S2**). Due to the wide range of studied habitats, the parasite list comprised over 30 species, many of which were reported only once. Thus, we selected only the most frequent for the analysis, recorded in more than 50% of locations.

### 2.4 Statistical analysis

Metazoan parasites tend to be aggregated and overdispersed within the host population [1, 38, 39], resulting in overdispersion or an excess of zeros in the data. However, many studies modelling parasite load in relation to environmental variables have not accounted for these two factors. Consequently, the misspecified models may have produced unreliable parameter estimates and distorted test results.

Here, we overcame this drawback by using zero-inflated models implemented in the glmmTMB R package [40]. The zero-inflated models assume that zero, i.e., the absence of a parasite in a host, results from a combination of two biologically different processes: 1) a host is not infected because it did not encounter the infective stage of a parasite (the parasite was either absent or at a very low level in the environment), or 2) a host encountered the parasite but did not develop an infection, e.g., due to immunity. Biologically, such an approach allows accounting for the differences between analysed sites or seasons in the risk of infection that may be associated with altered survival of the infective stages, lower transmission risk between hosts, etc. Mathematically, it provides a better-fitting model compared to other methods that takes into account the overdispersion and/or excess zeros.

In the analysis at the temporal and medium-spatial scales, we focused on individual levels of infection, rather than per-population or per-site means, as suggested by [31], who pointed out that the outcome of a model may be misleading when several functional groups of voles are used for calculating the overall prevalence. Moreover, studies examining the relationship between infection intensity and climatic conditions are missing [16]. Thus, the response variable in the models was parasite loads per individual. The prevalence (percentage of infected individuals per location) was fitted as a response variable only in the models including the large-scale data where infection intensity was not available, since the individual-level infection data were not reported in the published studies.

First, we tested for overdispersion and excess of zeros using DHARMa [41], and if both turned significant, we fitted zero-inflated models with negative binomial errors (ZINB). As the response variable, we used parasite species that infected more than 5% of voles, building separate models for each parasite species and each dataset (spatial or temporal). For the large-scale spatial dataset based on published data, where the individual parasite burdens were not available, we used the percentage of infected hosts per location, and we fitted the glmmTMB function with Beta errors.

In each model, as explanatory variables we fitted host sex, host body mass, and population density as conditional parameters, and climatic variables were fitted in the conditional and zero-inflated parts. Although density may affect the likelihood of infection through increased deposition of infected faeces at high densities, it also affects the host physiology through increased stress, increased competition, lower resources per capita, etc. Since it is not possible to disentangle these two types of effects, we decided to keep this variable only in the conditional part of the model to have a clear picture of the effect of the weather conditions.

Climatic variables are often correlated (e.g., snow cover correlates with temperature), and collinearity among explanatory variables increases the uncertainty of the parameter estimates and distorts model interpretation. Thus, based on the correlation matrix **(Tables S4 and S5**), for the models, we selected uncorrelated variables that potentially had a major impact on parasite egg survival in the environment [42]. In the temporal model, these were the average temperature at the ground and the precipitation. In the spatial models, we fitted either precipitation or soil humidity, and we selected the model with a better fit based on the lower AIC. Additionally, in the models we included latitude expressed in decimal degrees. Percentage of the explained variance (R^2^) of the model was calculated using the performance package [43] with a function dedicated to zero-inflated models. The effects of model parameters were interpreted and visualised using the ggpredict function from the ggeffects package [44]. The function produces adjusted predictions (model-based estimates) of the response variable for different combinations of predictor values, which allows for inference of the expected outcome for given values of the predictors.

## 3. Results

### 3.1. Climatic conditions affecting parasite load over time

The most prevalent helminth species in the temporal dataset was the nematode *Heligmosomum mixtum,* infecting 64.3% of voles overall, followed by *Aspiculuris tianjinensis* (20.3%) and the tapeworm *Catenotaenia hentonneni* (17%). The remaining species were less frequent: the nematode *Mastophorus muris* was present in 9.8% of voles, *Aonchotheca annulosa* in 9.5%, and the cestode *Paranoplocephala omphalode*s in 5%. Other cestode species were less frequent in voles, with a collective prevalence of 5.3%. *Syphacia sp*. and *Heligmosomoides glareoli* were rare, present in only 3.2% and 1.3%, respectively.

In the temporal dataset, we found considerable variations in temperatures between years. The average ground temperature in September varied from 12.2 to 15.6°C, and in May from 11 to 13.9°C (**Figure 1**). There was an apparent increasing trend in temperatures, confirming the overall rise of temperatures over the past decades, yet in the case of our data, it was not statistically significant (LM, May β = 0.1415, p = 0.149; September β = 0.1591, p = 0.118).

**Figure 1.**
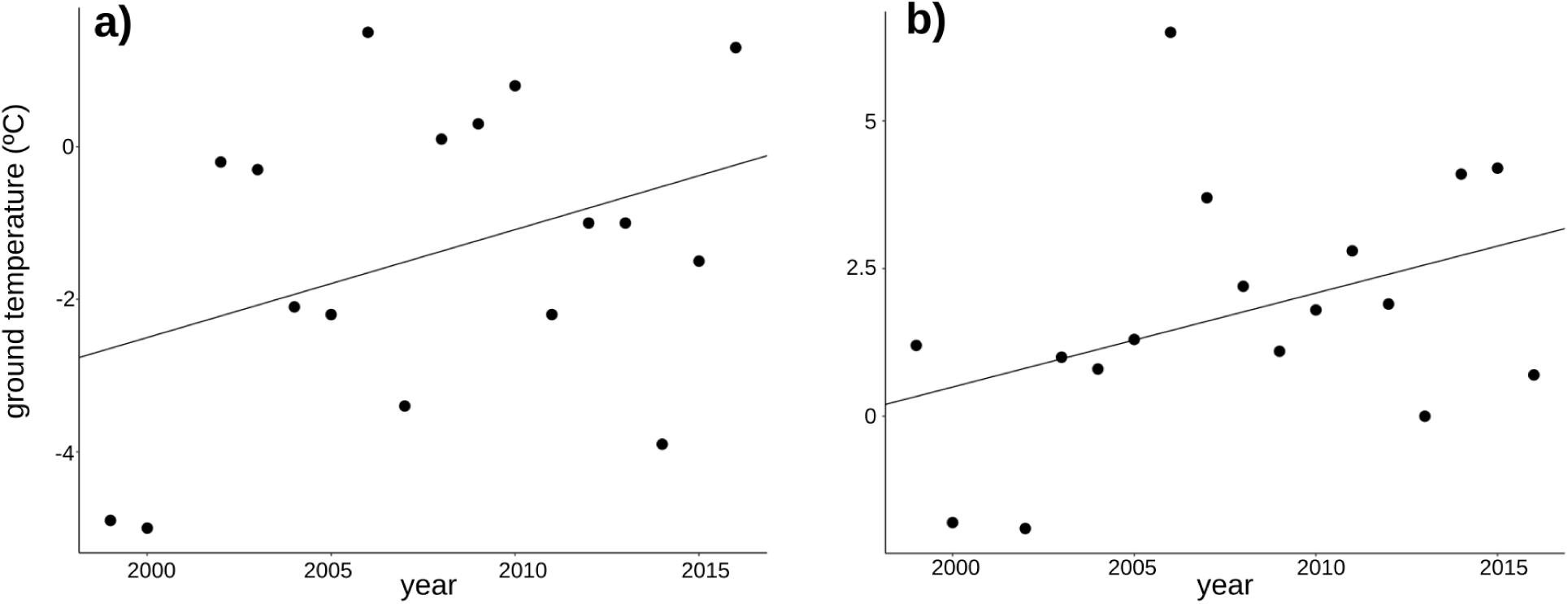
Year-to-year variation in the ground temperatures in a) May and b) September at the site of the temporal dataset.

Temperature and precipitation 2 months prior to sampling had a strong impact on the infection with *the most frequent nematode H. mixtum* (β_condtional_= −0.062, p < 0.0001 for temperature, β_condtional_= −0.09, p < 0.0001 for precipitation, **Table 1**). Generally, the higher the temperature, the lower the parasite burden of the infected individuals, and the effect was much stronger at negative temperatures. It is important to note that the voles were not captured in winter, and the extra low temperatures denote cold early spring (March), two months before the first sampling session in May. Change from −20°C to −3.9°C decreased the parasite load from 10.8 to 3.9 worms per infected vole, yet due to the low number of months with ground temperatures below 10°C, the estimates had low precision (**Figure 2a**). The high precipitation affected *H. mixtum* burdens negatively, yet the effect was not as strong as in the case of temperature: an increase from 10 to 16 days with precipitation decreased the parasite load by almost one worm (from 2.99 to 2.09) in the infected individuals (**Figure 2b**). Contrary to predictions, neither temperature nor precipitation affected the probability of encountering the parasite, as indicated by the non-significant effects in the zero-inflated parts of the model. The only exception was the marginally significant effect of temperature in the zero-inflated part of the model of infection with *H. mixtum* at time 0.

**Figure 2.**
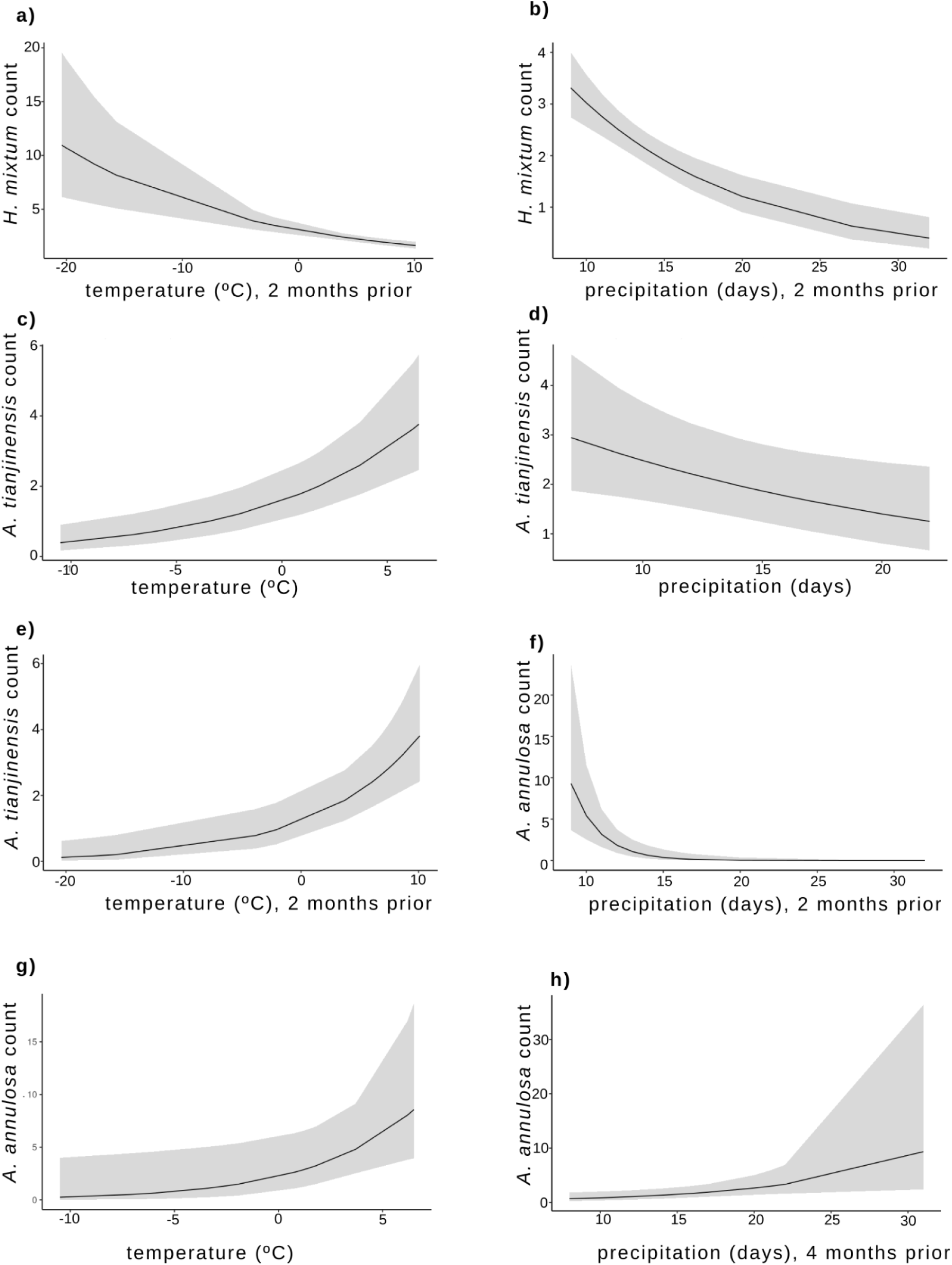
Predicted (conditional) parasite counts in relation to ground temperature and/or precipitation. Only significant effects are shown. Predictions from models with C. *hentonneni* or *M. muris* as the response variable are not shown due to the low explanatory power of these models (R^2^<0.1)

**Table 1.**
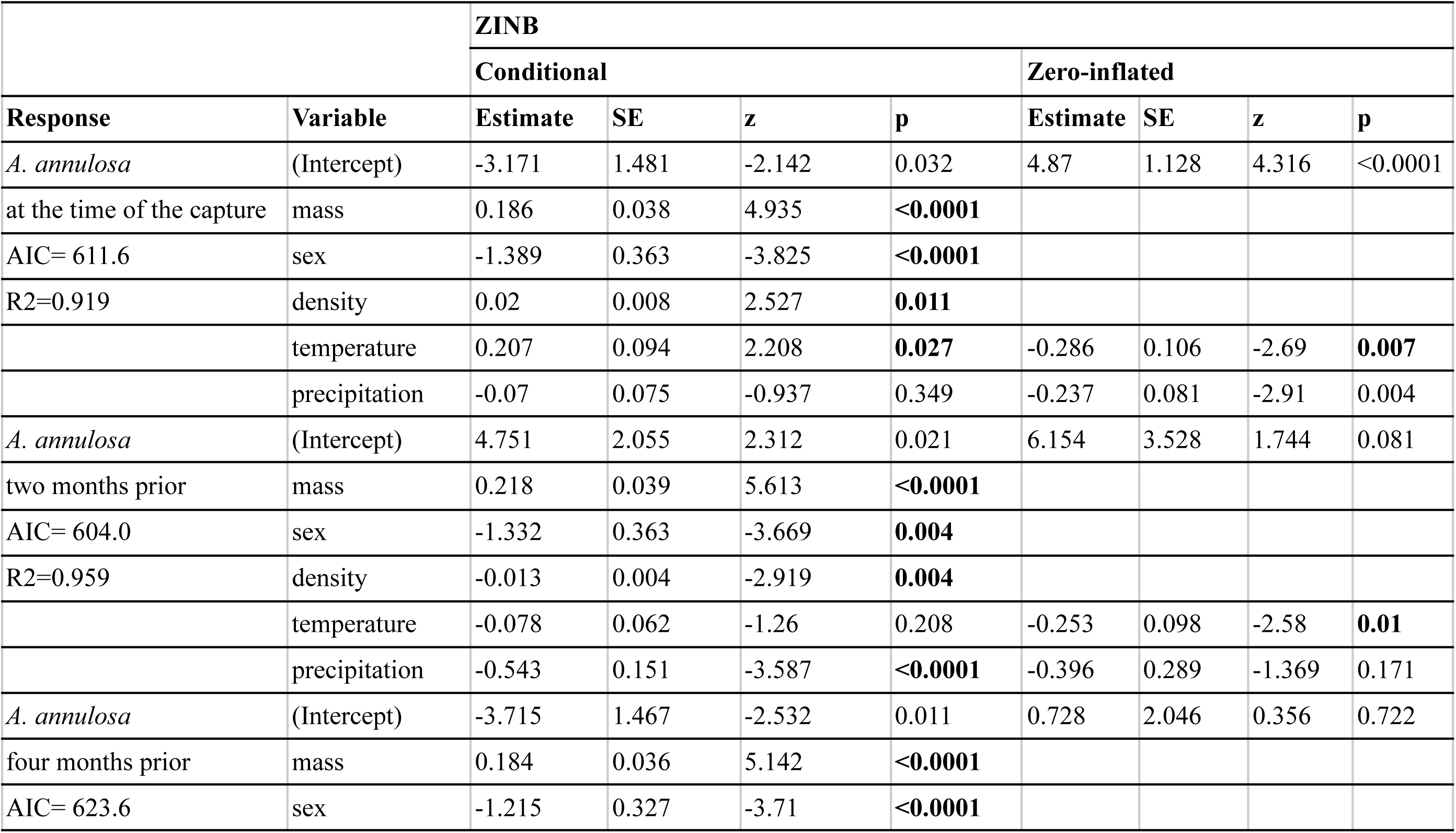
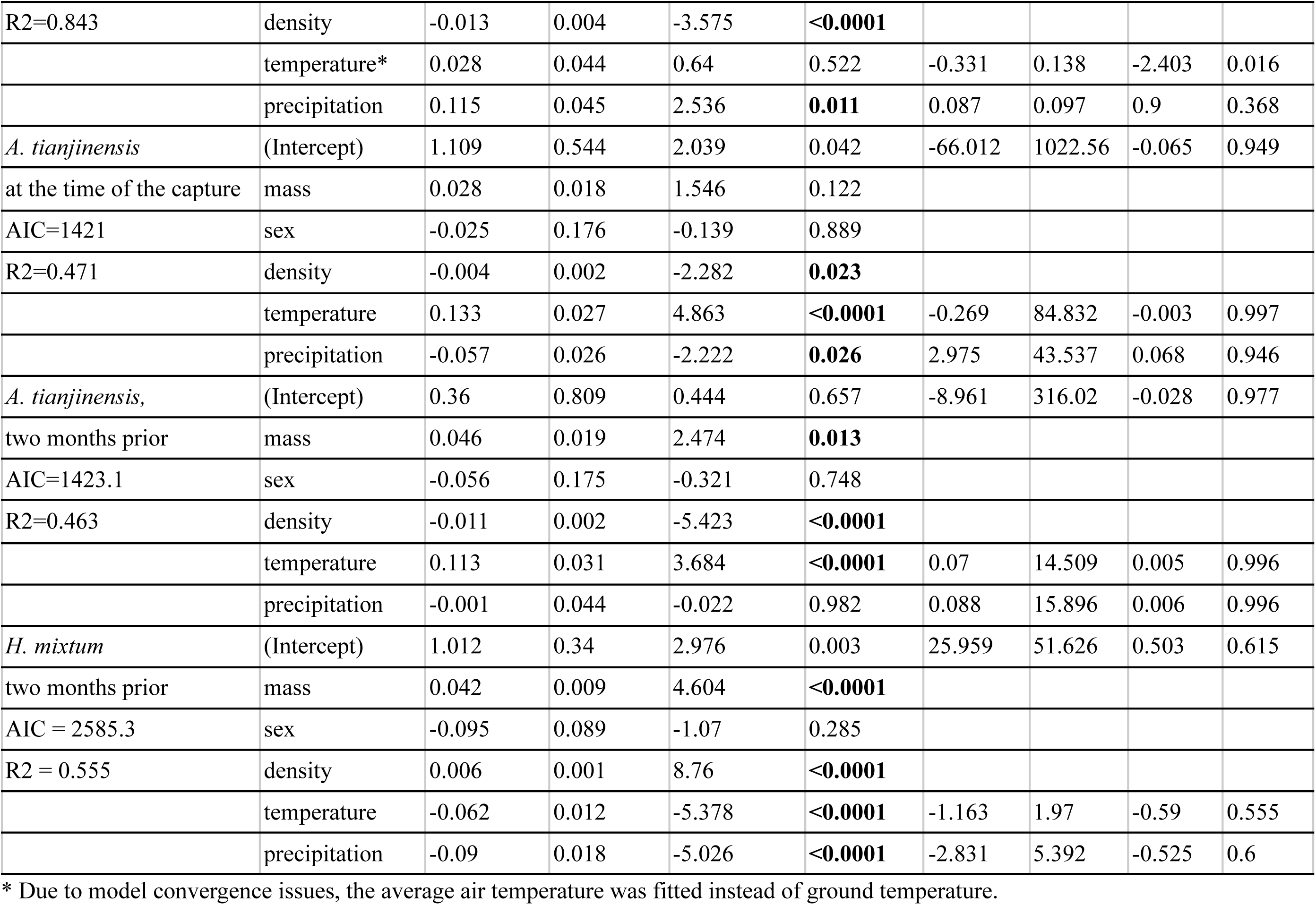
Summary of significant zero-inflated negative-binomial models showing the effects of weather conditions on parasite counts over time. Separate models were constructed to account for the weather conditions at the time of the capture, two, and four months prior what is indicated next to the species name. Explanatory variables: mass - host body mass, sex - host sex, density - host population density at the time of sampling, temperature - ground temperature in a given month, precipitation - number of days with rainfall or snowfall in a given month. R2 is the amount of variance explained by the model. Full model are presented in the Table S6.

**Table 2.** Summary of significant zero-inflated negative-binomial models showing the effects of weather conditions on parasite counts across medium spatial scale. Explanatory variables: mass - host body mass, sex - host sex, density - host population density at the time of sampling, temperature - ground temperature in a given month, precipitation - number of days with rainfall or snowfall in a given month. R2 is the amount of variance explained by the model. Full model outputs are presented in the Table S7.

|  |  | **ZINB** |  |  |  |  |  |  |  |
| --- | --- | --- | --- | --- | --- | --- | --- | --- | --- |
|  |  | **Conditional** |  |  |  | **Zero-inflated** |  |  |  |
| **Response** | **Variable** | **Estimate** | **SE** | **z** | **p** | **Estimate** | **SE** | **z** | **p** |
| *A. annulosa* | (Intercept) | 27.824 | 9.825 | 2.832 | **0.005** | 34.47 | 75.98 | 0.454 | 0.65 |
| AIC=365.3 | mass | 0.21 | 0.036 | 5.797 | **<0.0001** |  |  |  |  |
| R2=0.815 | sex | -1.238 | 0.39 | -3.174 | **0.002** |  |  |  |  |
|  | density | 0.001 | 0.006 | 0.085 | 0.932 |  |  |  |  |
|  | latitude | -0.083 | 0.122 | -0.683 | 0.495 | -0.455 | 1.056 | -0.43 | 0.667 |
|  | temperature | -3.022 | 0.689 | -4.384 | **<0.0001** | -3.673 | 2.02 | -1.818 | 0.069 |
| *H. mixtum* | (Intercept) | -48.94 | 13.568 | -3.607 | **<0.0001** | 100.014 | 66.346 | 1.508 | 0.132 |
| AIC= 647.8 | mass | 0.098 | 0.021 | 4.575 | **<0.0001** |  |  |  |  |
| R2=0.241 | sex | 0.184 | 0.215 | 0.859 | 0.39 |  |  |  |  |
|  | density | -0.003 | 0.003 | -1.001 | 0.317 |  |  |  |  |
|  | latitude | 0.406 | 0.178 | 2.276 | **0.023** | -2.973 | 1.952 | -1.523 | 0.128 |
|  | temperature | 2.226 | 0.473 | 4.703 | **<0.0001** | 4.715 | 4.689 | 1.006 | 0.315 |
|  | precipitation | 0.01 | 0.002 | 4.906 | **<0.0001** | 0.008 | 0.009 | 0.957 | 0.339 |

Temperature played an important role in *A. tianjinensis* infections. Unlike in the case of *H. mixtum*, the number of worms in an infected host increased with the temperature at time 0 and time −2 (β_condtional_ = 0.027, p < 0.0001 and β_condtional_= 0.031, p < 0.0001 respectively, **Figure 2c** and **2d**). There was no effect of soil humidity on *A. tianjinensis* infections.

Temperature had a significant effect on infections with *A. annulosa* (**Figure 2e-h)**. Interestingly, in the zero-inflated part of the models with *A. annulosa* at time 0 and −4 the effect was negative (β_zi_ = −0.286, p = 0.007 and −4 β_zi_ = −0.286, p = 0.016 respectively) but positive in the conditional part for time 0 (β_condtional_ = 0.094, p = 0.027). This suggests that the odds of being infected decrease with temperature, but once infected, the host has a higher parasite load at higher temperatures. The more days with precipitation, the higher parasite load at time −2 (β_condtional_= −0,543, p = 0.0004), but the effect was opposite, although much weaker and associated with higher uncertainty, at time −4 (β_condtional_= 0.115, p = 0.011). Notably, models for *A. annulosa* had a very high fit, explaining >90% of variance.

In all models, we found a significant effect of host density on parasite load, with higher infection rates at higher population densities. Density may negatively affect host physiology at high values, but it can also be linked to a higher density of parasite transmission stages (embryonated eggs and infective larvae) on the ground. Hence, we re-ran the models with density fitted also in the zero-inflated part, and it did not affect the significant effects of the weather conditions (results not shown).

Models for *M. muris* and the cestodes *C. hentonneni* and *P. omphalodes* had a very low fit, explaining less than 10% of the variance. Thus, it is not possible to make a meaningful prediction based on the resulting estimates. Full models for all analysed parasites are given in the Table S6.

### 3.2. Climatic conditions affecting parasite load on a medium geographic scale

Similar to the temporal dataset, *H. mixtum* was the most prevalent parasite, infecting 33.7% of voles across 16 locations. Cestodes were found in 22.2% of voles. *A. tianjinensis* infected 21.15%, *M. muris* 12.2%, and *A. annulosa* was found in 11.8%. *Syphacia sp*. and *H. glareoli* were rare, present in less than 2.5% of hosts.

We did not find an effect of climatic conditions on the zero-inflated part of the models, which suggests that the likelihood of infection was similar across locations, regardless of the climatic differences. Contrary to our predictions, latitude generally did not significantly affect parasite burdens, except for *H. mixtum*. Voles from the southern part of the analysed range had fewer *H. mixtum* than those collected further north (β_condtional_ = 0.406, p = 0.023, **Figure 3**). This effect cannot be attributed to any differences in host densities, as there was no correlation between density and latitude (r = 0.0191, p = 0.751). Moreover, the southernmost populations (latitude below 50°N) were located in two regions with contrasting population densities - in Křižanovská vrchovina host density was the highest, reaching 140 voles per 10^-4^ trap-hours while in Beskid Żywiecki densities were low (9-30 voles per 10^-4^ trap-hours). In the model for *H. mixtum*, we also found a strong and significant effect of the ground temperature and precipitation (**Figure 3**). Parasite load was higher in locations with higher annual mean temperatures and higher precipitation.

**Figure 3.**
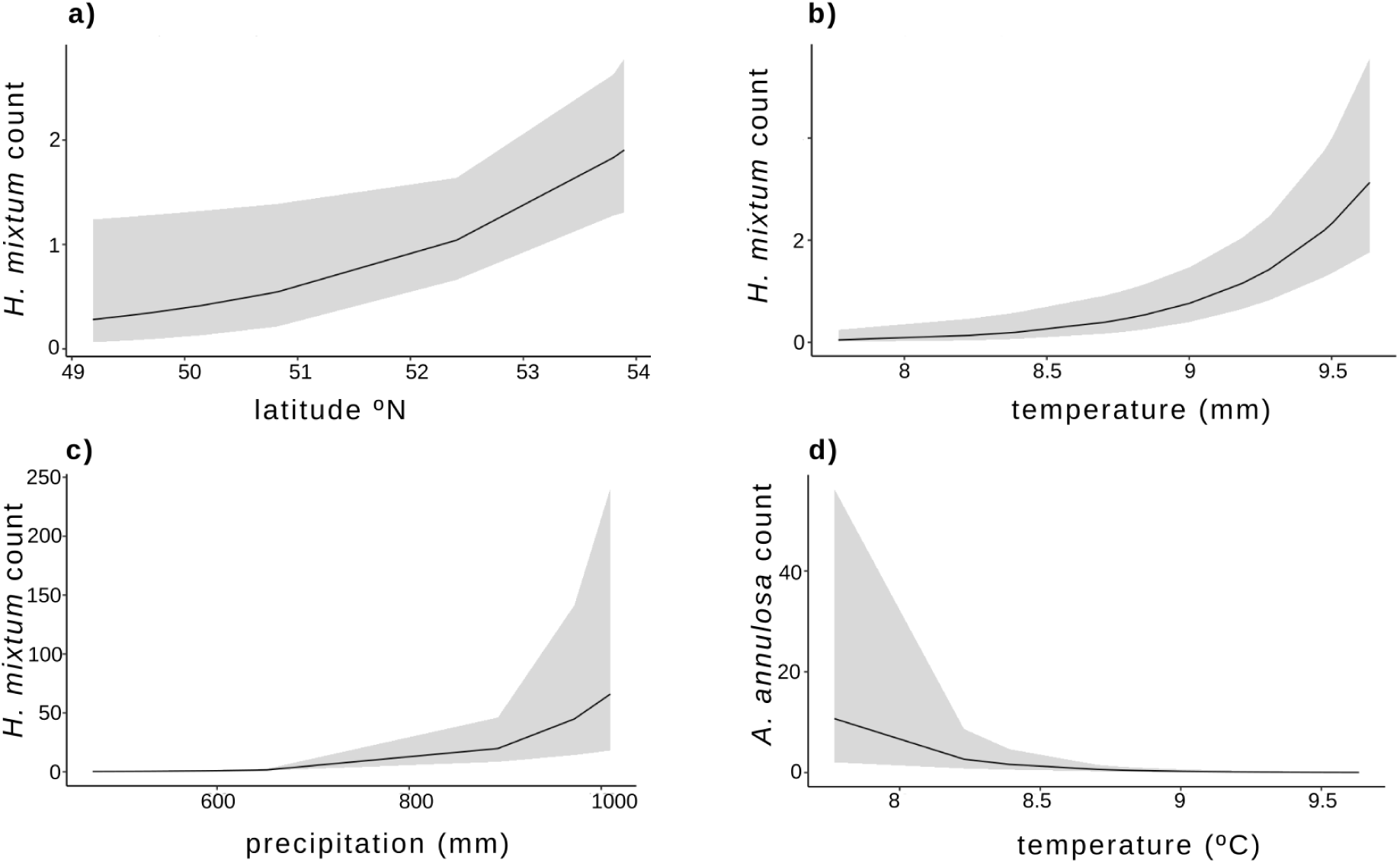
Predicted parasite counts in relation to annual mean ground temperature and/or precipitation at the medium spatial scale.

Soil humidity affected cestode infections, with parasite burdens decreasing as humidity increased, yet the model had very low explanatory power (R² = 0.030), so this result should be interpreted with caution. Climatic conditions did not affect infections with *A. tianjinensis* or *M. muris*. We found a strong effect of temperature on the intensity of infections with *A. annulosa*, and the model explained over 80% of the variance in the data. The highest parasite loads were found in locations with the lowest ground temperatures, yet the effect was only visible for temperatures below 8°C, while for the highest temperatures, there was no effect (Figure 2). Full models for all analysed parasites are given in Supplementary Materials, Table S7.

### 3.3. Climatic conditions affecting parasite prevalence at a large geographic scale

Overall, the authors reported 53 helminth species infecting bank voles, including 23 nematodes, 23 cestodes, and 7 flukes (**Table S8**). The most prevalent species were *H. mixtum* (reported in 57.1% of the studied locations), *H. glareoli* (50%), *A. annulosa* (42.8%), *M. muris* (42.8%), and *Syphacia petrusewiczi* (39.2%). The most prevalent cestode was *C. henttoneni* (39.3%). Trematodes were observed sporadically. Almost half of the species (24) were reported from a single location. In the models, we considered only those species that were reported in more than 10% of the locations.

The only models where we found a significant effect of climatic conditions included *M. muris*, which was more frequent at sites with lower precipitation (**Table 3**, **Figure 4b**), and the cestode *C. hentonneni,* which was more often reported from locations with low precipitation and low temperatures (**Figures 4a-b**).

**Figure 4.**
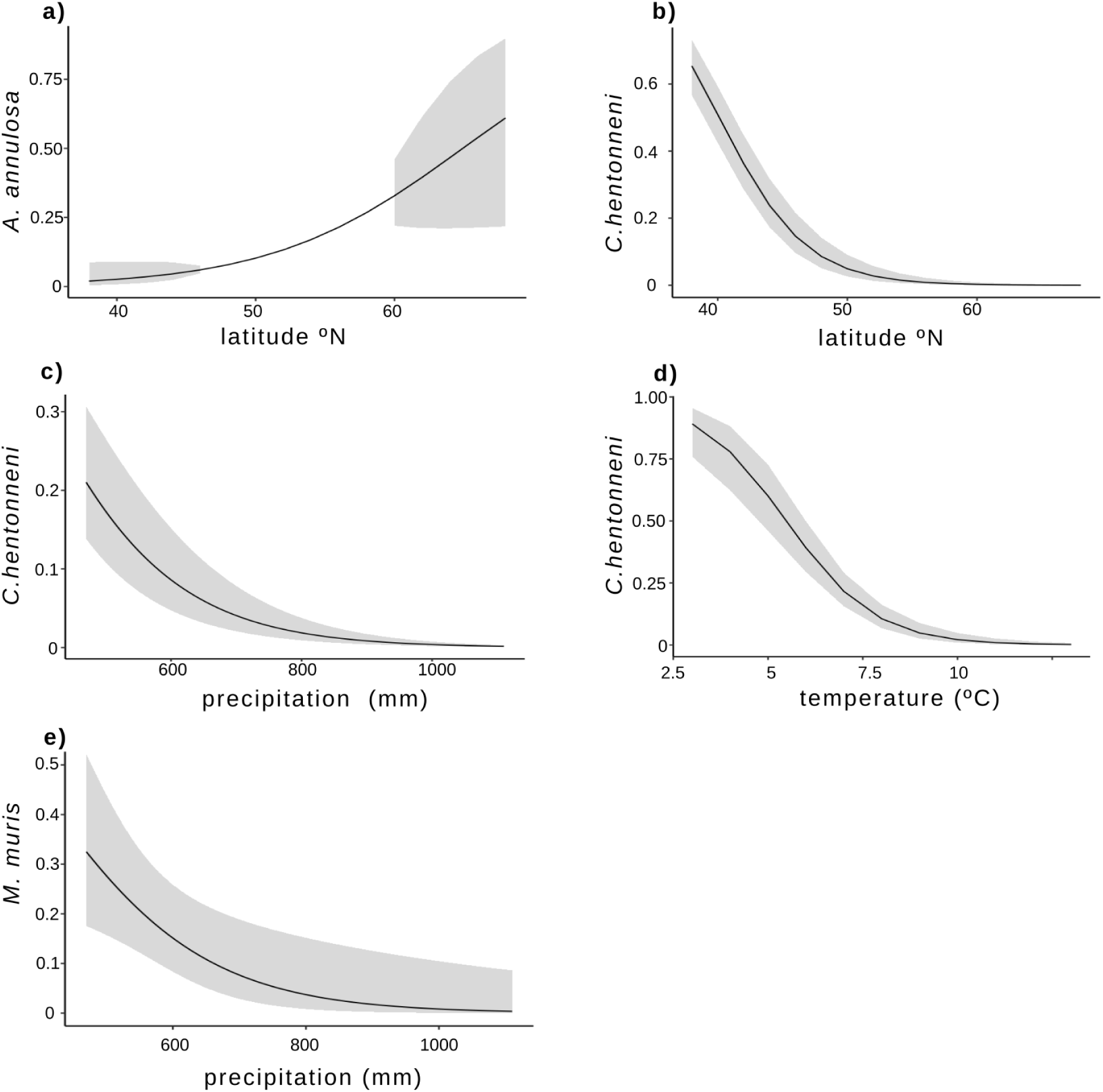
Predicted parasite prevalence in relation to annual mean ground temperature and/or precipitation across the large spatial scale.

**Table 3.** Summary of significant zero-inflated negative binomial models showing the effects of weather conditions on per-site parasite prevalence across the large spatial scale. Explanatory variables: mass - host body mass, sex - host sex, density - host population density at the time of sampling, temperature - ground temperature in a given month, precipitation - number of days with rainfall or snowfall in a given month. R2 is the amount of variance explained by the model. Full model outputs are presented in the Table S9.

|  |  | **ZINB** |  |  |  |  |  |  |  |
| --- | --- | --- | --- | --- | --- | --- | --- | --- | --- |
|  |  | **Conditional** |  |  |  | **Zero-inflated** |  |  |  |
| **Response** | **Variable** | **Estimate** | **SE** | **z** | **p** | **Estimate** | **SE** | **z** | **p** |
| *C. henttoneni* | (Intercept) | 25.3642 | 4.7559 | 5.333 | **0** | -7.6809 | 14.3826 | -0.534 | 0.5933 |
| AIC=-12.7 | n | 0.0037 | 0.0015 | 2.528 | **0.0115** |  |  |  |  |
| R2=0.961 | latitude | -0.3005 | 0.0472 | -6.369 | **0** | 0.1986 | 0.1891 | 1.05 | 0.2936 |
|  | precipitation | -0.008 | 0.0022 | -3.552 | **0.0004** | -0.0098 | 0.0047 | -2.066 | **0.0388** |
|  | temperature | -0.8491 | 0.1363 | -6.23 | **0** | 0.5673 | 0.5586 | 1.016 | 0.3098 |
| *M. muris* | (Intercept) | 7.4712 | 8.1858 | 0.913 | 0.3614 | -7.8575 | 9.0389 | -0.869 | 0.385 |
| AIC=24.4 | n | 0.0019 | 0.0022 | 0.889 | 0.3741 |  |  |  |  |
| R2=0.385 | latitude | -0.1024 | 0.1073 | -0.955 | 0.3398 | 0.0903 | 0.1124 | 0.804 | 0.421 |
|  | precipitation | -0.0076 | 0.003 | -2.579 | **0.0099** | 0.003 | 0.0026 | 1.17 | 0.242 |
|  | temperature | 0.056 | 0.3681 | 0.152 | 0.8791 | 0.169 | 0.324 | 0.522 | 0.602 |

## 4. Discussion

In this study, we examined the effects of climatic variables (temperature and humidity or precipitation) on parasite burdens in a wild rodent across temporal and spatial scales with focus on individual parasite species and individual parasite burdens per host rather than parasite communities and population means as used in previous studies. As pointed out recently [16], the studies on the relationship between infection intensity and climatic conditions are rare, so our study fills this gap.

Our results show the effect of climatic conditions was visible at different scales and that climate-parasites relationship varies depending on the scales and the parasite species. For instance, in the temporal scale, higher parasite loads with *H. mixtum* were observed at lower temperatures (regardless of the season), but in the medium-spatial scale, it was higher in locations with higher annual temperatures. The main reasons for the observed discrepancies across species and scales are likely to stem from differences in parasite biology, but also from differences between locations in parasite community compositions. Thus, our results underline that depending on the scale of the study, the outcome may be strikingly different. This finding should be taken into account when constructing predictions about parasite occurrence.

### Temporal scale: weather and parasite survival

Despite numerous studies analysing temporal variation in parasite burdens in rodents, most have simply reported associations between infection levels and trapping sessions [e.g., 25]. However, in temperate zones, mean monthly temperatures can vary substantially from year-to-year, as shown in Figure 1. This emphasises the importance of including actual temperature data rather than proxies such as “season” or “month” in statistical models. Here, we accounted not only for seasonal variation but also for weather conditions two and four months prior to trapping, reflecting the time required for parasite eggs and larvae to develop into infective stages and for hosts to acquire infections.

Significant effects of ground temperature were observed in the zero-inflated component of models for all nematode species studied, indicating that the likelihood of encountering parasites varies with temperature. Once parasites were present in the environment, their burdens were further influenced by environmental conditions, as reflected in the conditional component of the models. Temperature effects were species-specific: while *H. mixtum* burdens decreased with increasing temperature, burdens of *A. tianjinensis* and *A. annulosa* increased. Models predictions (Figure 2) suggest that milder winters, particularly fewer months with average temperatures below 0**°**C, are expected to have a major effect on nematode infections in the bank voles.

High *H. mixtum* burden at low temperatures is a good example of the confounding effects of weather and host dynamics [31]. In spring, captured animals belong to the previous autumn cohort that survived winter. These older individuals (>6 months) have had time to accumulate parasites over their lifetimes, and they are the only group exposed to negative temperatures, explaining the observed association between cold conditions and high parasite loads. In contrast to temperatures, precipitation had a more uniform effect across species, parasite burdens decreased with the number of rainy or snowy days. This is consistent with the findings of [45], who observed that humidity and temperature influence larval migration from the faeces to grass, affecting host exposure.

The importance of climatic conditions in explaining the seasonal dynamics of helminth infections has been reported earlier, in particular in the case of *H. mixtum* [46] in voles from Finland. They found the highest prevalence of *H. mixtum* after cold and rainy summers. Similarly, in our study, low temperature two months prior to sampling was associated with higher parasite load, but high precipitation resulted in decreased parasite burdens. The discrepancy between [46] and our findings may be attributed to the fact that [46] studied cyclic populations inhabiting the northern boreal zone, compared to the temperate continental zone analysed in the current study.

All three species showing significant effects in this study have direct life cycles. Laboratory studies of *A. tianjinensis*’s congener, *A. tetraptera*, indicate that adults survive up to 50 days in the proximal colon of mice [47], comparable to post-weaning survival rates of hosts [48]. Eggs expelled in feces require incubation in soil (6-7 days at 24°C) to become infective [49]. They are resistant to environmental stresses, with oxygen, temperature, and humidity all essential for development [49,50]. Optimal development occurs between 20-30°C, whereas 0-4°C prevents egg development and 37°C reduces infective egg yield [49].

*H. mixtum* females release eggs that hatch into first-stage larvae (L1), progressing through L2 and L3. While environmental survival of *H. mixtum* larvae has not been studied directly, related heligmosomids (*Heligmosomoides bakeri* = *Nematospiroides dubius*) show that free-living larvae survive various environmental conditions until ingested by a host. Eggs hatch at 20°C within 34–36 h, and at 23–28°C within 26 h; L3 larvae develop in 4–6 days and can remain dormant for extended periods. Larvae retain the L2 cuticle as a protective sheath until ingestion by a host [51, 52].

The genera *Aspiculuris* and heligmosomids differ in environmental strategies: *Aspiculuris* larvae remain protected within eggs until ingestion, with specialized layers providing thermal tolerance and resilience to environmental extremes [53, 54]. This likely explains why *A. tianjinensis* burdens increased with temperature, whereas *H. mixtum* burdens decreased. Heligmosomid larvae hatch in the environment, making them more vulnerable to high temperatures and environmental stressors [55, 56].

### Spatial scale: local (in)stability

The spatial scale models present different resolutions compared to temporal models. While temporal analyses consider month-to-month variation, spatial models are based on multi-year average temperatures and precipitation. Moreover, in the large-scale model including published data, we used the percent of infected hosts per population instead of individual parasite loads to account for aggregated patterns. Species-specific responses to climate were evident. The parasite burden of *H. mixtum* increased at temperatures above 9°C, whereas *A. annulosa* burdens decreased sharply above 8.5°C. Higher precipitation (>800 mm) was associated with higher *H. mixtum* and lower prevalence of *C. hentonneni* and *M. muris,* likely reflecting the distribution of intermediate invertebrate hosts and their climatic niches.

Interestingly, at the largest geographic scale, no significant association was observed between the most prevalent nematode, *H. mixtum,* and climatic conditions, despite clear relationships at temporal and medium spatial scales. Similarly, recent metanalysis of the relationship between infection and climatic conditions in the terrestrial animals found large variation in the effect of temperature and precipitation on parasite prevalence, and temperature on infection intensity [16]. We hypothesise that this pattern likely reflects the reduced resolution that occurs when multiple local populations or sampling seasons are aggregated. The drawbacks of such an approach have been thoroughly discussed by [31], who emphasized the importance of considering separate functional groups of rodents, e.g., subadults or overwintered, as parasite burden in these groups may differ considerably even within the same season. By using per-individual parasite loads, our models avoided this limitation. Similarly, [14] reported that remotely sensed vegetation indices explained approximately 30% of within-habitat variation in overall infection burden in voles but noted that this approach is less reliable for comparisons between localities. These findings highlight the importance of considering both spatial resolution and host cohort structure when interpreting parasite-environment relationships.

### Effect of host population dynamics and predicted impacts

Climate change, particularly increasing temperatures and milder winters, may allow rodents to begin reproducing earlier in spring or remain reproductively active throughout the winter [57]. These changes could alter host population dynamics and, consequently, parasite dynamics. However, the direction, magnitude, and effects on parasite loads remain difficult to predict [58].

Previous studies on the population dynamics of vole helminths showed that the density of the host species (*M. glareolus*) does not directly influence infection levels of common helminths such as *H. mixtum* and *Catenotaenia sp*. Nonetheless, fluctuations in the host population density led to corresponding changes in the prevalence of these common helminths [59]. In contrast, rare helminth species, including *M. muris*, *A. annulosa*, *Sphacia petrusewiczi*, *Paranoplocephala kalelai*) exhibit less predictable dynamics and are less affected by low host density [21, 23, 59].

Interactions among intestinal helminths species are also influenced by the host’s immune responses, though these interactions explained only a small proportion of the variation in abundance [21]. Both immunological and hormonal responses play important roles in regulating helminth populations. For common helminths, host immune regulation can strongly influence population dynamics, with transmission notably impaired only at extremely low host densities. Rare helminths, however, are more affected by stochastic survival during infective stages and by seasonal variation in transmission [60, 61]. The cumulative effect of such within-host regulation can be more pronounced at the population level, influencing multi-year transmission dynamics, especially under fluctuating host densities and changing climatic conditions that affect host physiology and immune function.

Life history traits of helminths, such as feeding mode, reproduction, and life cycle stage, do not appear to regulate population dynamics [61]. Similarly, geographic studies of *M. glareolus* helminth assemblages show that life cycle types do not determine seasonal or long-term variations in helminth populations. Across continental Europe, the helminth fauna of *M. glareolus* is highly predictable, with patchy spatial distribution and strong aggregation, reflecting differences in individual host exposure and susceptibility rather than large-scale environmental heterogeneity [24, 62]. These findings highlight the role of host ecology and behaviour, which can be strongly influenced by climatic variation, in shaping spatial and temporal patterns of parasite infection.

Population fluctuations in small rodents correlates positively with latitude and snow cover, factors that also influence the abundance of alternative prey for generalist predators. Areas with deeper snow cover, at the same latitude in Eastern Europe show higher cyclicity indices compared to regions with poor snow cover, such as Great Britain and Fennoscandia [63]. This predator-mediated regulation likely contributes to more stable microtine populations observed in southern Fennoscandia and central Europe [64, 65]. Since parasite transmission depends on available hosts acquiring infection, such latitude- and snow-related differences in rodent population stability are likely to translate into spatial variation in parasite prevalence and persistence under changing climatic conditions.

### General conclusions

In this study, we show that temperature and soil humidity/precipitation influence parasite burdens in small rodents in a way that strongly depends on temporal scales, spatial scales and parasite species. Importantly, this variation is not random but reflects underlying differences in parasite biology and local ecological conditions (Figure 5). Our results indicate that short-term climatic fluctuations (seasonal variation) can play a stronger role in shaping local host-parasite dynamics than consistent year-to-year changes. In contrast, broader spatial patterns are driven by environmental heterogeneity and differences in parasite communities composition among locations, as seen in high between-site variation observed, even in locations with similar climatic conditions. It also seems that the parasite life cycle may play a role: while species with direct life cycle, where infective eggs persist in a soil, are more susceptible to temporal variation in the weather conditions, species with the complex life cycle did not show such a pattern. Instead, in those we found large-scale associations with climatic conditions what suggest a crucial role of the range and distribution of the intermediate, usually invertebrate hosts.

**Figure 5.**
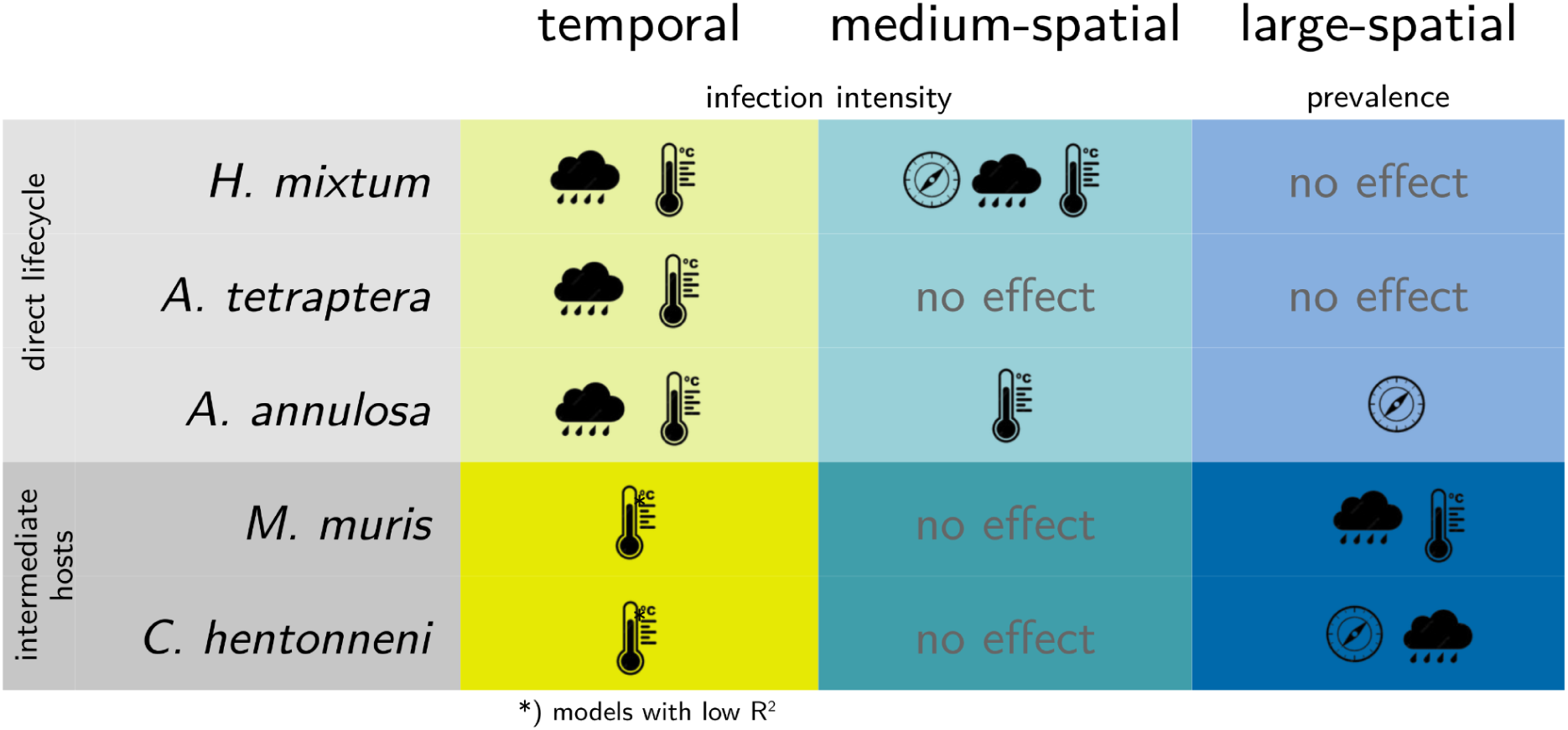
Synthetic summary of the effects studied in the current work.

These findings highlight that climate-parasite relationships are scale-dependent, and that predicting parasite dynamics under ongoing climate change require consideration of both spatial and temporal context. Rather than expecting uniform responses across parasite species, future studies should focus on identifying the ecological conditions under which climate effects are strongest.

## Authors contributions

AK conceived the ideas; AEO and AK acquired funding; UZP, JB, AB and AK, designed methodology; AEO, VIA, AD, AR, MA, DDS, KT, MG, JBB, JB, AB and AK collected the data; AEO, UZP and AK analysed the data; AEO and AK led the writing of the manuscript. All authors gave final approval for publication.

## Acknowledgements

We thank Joelle Gouy de Bellocq and Alena Fornůsková from the Czech Academy of Sciences, Institute of Vertebrate Biology, for their help in data collection in the Czech Republic. This work was supported by funding from the Polish National Science Foundation (NCN) grant number 2020/39/O/NZ8/01596 to AK and from the Polish National Agency for Academic Exchange (NAWA) grant number BPN/PRE/2022/1/00025/U/00001 to AEO.

