## Supplementary Figures S1- S2 for "Species-specific responses of helminths to temperature and moisture: long-term and multi-scale analyses in a free-living rodent"

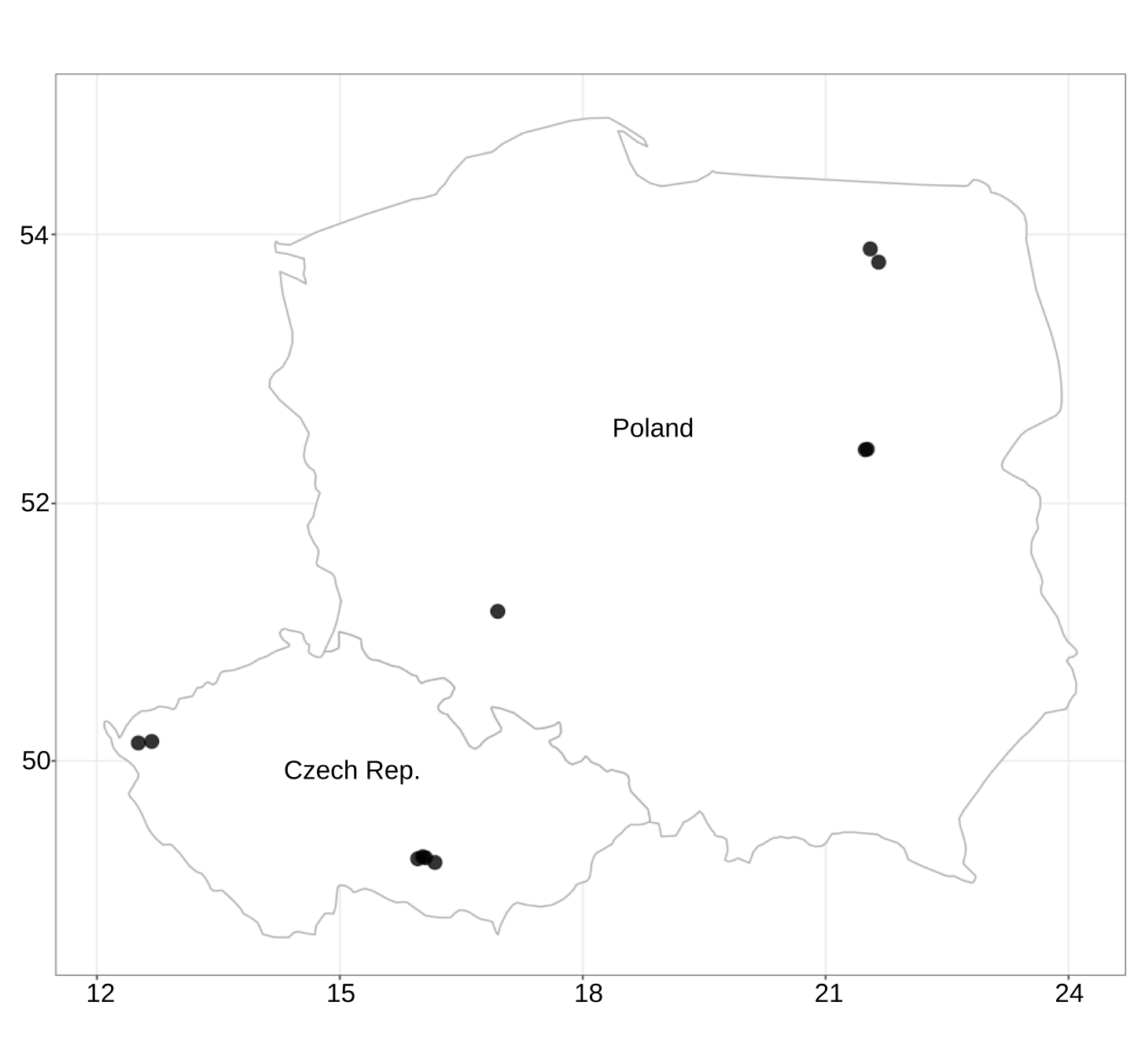
Figure S1. Location of data points in the medium-scale spatial set.

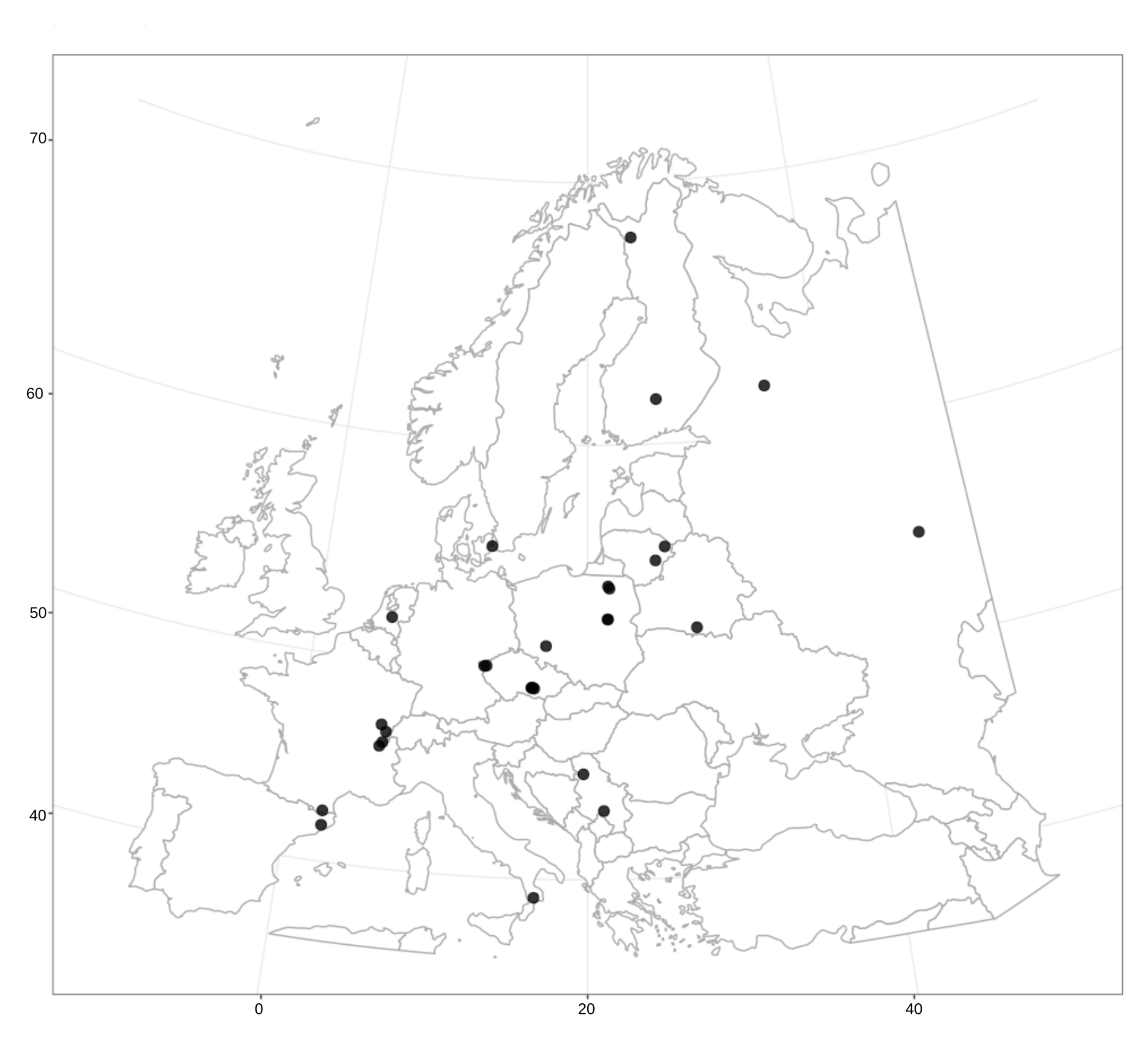

Figure S2. Location of data points in the large-scale spatial set.
